# On the influence of stratification and lake size on pelagic and benthic algal dynamics: A modelling study using ‘Lake2D’

**DOI:** 10.1101/2025.11.17.688809

**Authors:** Hugo Harlin, Karl Larsson, Åke Brännström, Sebastian Diehl

## Abstract

This study investigates the influence of mixed layer depth and lake size on the dynamics of pelagic and benthic algae in stratified lakes using a two-dimensional, spatially explicit modeling approach. To this end, we modified the process-based, two-dimensional model ‘Lake2D’ to account for stratification in lakes by simulating a shallow layer with fast turbulent mixing, and a deep layer with slow turbulent mixing, separated by a sharp boundary. Using this modified version of Lake2D we comprehensively explored pelagic and benthic algal dynamics across a wide range of ecological conditions by varying lake mean depth, total nutrient content, lake size, and mixing depth in a full-factorial manner. Our results reveal distinct regimes of whole-lake algal dynamics: In shallow, nutrient-poor lakes, pelagic algae are nutrient-limited and sparse, while benthic algae thrive and constitute the majority of the total biomass, with no notable influence of mixed layer depth on algal dynamics. As lakes become deeper or more nutrient-rich, pelagic biomass increases and can become the dominant constituent of total biomass. For deep lakes, the model shows a characteristic L-shaped pattern in algal biomass for varying lake size and mixing depth. Pelagic algae exhibit low biomass in small lakes with a deep mixed layer, and high biomass otherwise. Benthic algae show the opposite pattern, exhibiting high biomass in small, deep lakes with a deep mixed layer, and significantly lower biomass otherwise. The study also uncovers a novel phenomenon, a benthic deep chlorophyll maximum, where the benthic algal biomass peaks at an intermediate depth along the lake bottom. This phenomenon arises in lakes of moderate depth and nutrient content, where a shallow mixed layer enables opposing vertical gradients of light and nutrients. These findings highlight the role of mixed layer depth and horizontal transport in structuring the spatial distribution and overall composition of algal biomass in lakes, underscoring the importance of vertical mixing and lake size for understanding lake primary production.

## 1 Introduction

Lake size, depth, and hypsography are fundamental physical characteristics that collectively have a profound influence on the abiotic conditions in lakes, and ultimately the ecological dynamics of algae therein that form the base of aquatic ecosystems [1–3]. In conjunction with the regional climate and material inputs from the catchment, these physical properties directly control key factors that regulate algal growth, such as light availability, nutrient cycling, and mixing regimes [4–6]. For example, in deep lakes, a greater water volume can dilute nutrient concentrations and reduce the light penetration to a small fraction of the water column, both of which tend to limit algal growth [7, 8]. Conversely, in shallow lakes, the constant circulation of water can keep dissolved nutrients accessible to algae, which, together with the favorable light conditions in these shallow systems, tends to support high levels of algal productivity [9].

In many lakes, yet another important physical property influencing algal growth and productivity is the depth of the mixed surface layer. Except for the most shallow water bodies, the majority of the world’s lakes are seasonally, intermittently, or permanently stratified into a surface layer with warmer, lighter water where turbulent mixing is high, and a layer with colder, denser water below where turbulent mixing is considerably lower [10–12]. In these lakes, the mixed surface layer is typically assumed to be the biologically most active part of the water column, to the extent that many theoretical and empirical studies of algal dynamics and productivity focus exclusively on patterns and processes within the mixed surface layer [13–16]. The role of mixing depth in algae-related processes has been explored extensively with mathematical models, which has led to the development and empirical testing of concepts such as the critical depth principle (predicting that algal blooms in deep waters bodies can only develop if the mixed surface layer is sufficiently shallow) [17–21], and the related prediction of a unimodal relationship between depth-integrated algal biomass and mixing depth [22, 23]. These studies often find a unimodal relationship between algal biomass and mixing depth [15, 23, 24]. A thorough understanding of the consequences of the mixing depth is critical because seasonal mixing regimes are changing with ongoing climate change [4, 12, 25–27]. A thorough understanding of the consequences of mixed layer depth for algal dynamics and productivity is therefore critical for predicting the effects of climate change in the future.

While the studies mentioned above have yielded many insights into the ecological consequences of mixed surface layer depth, they have focused exclusively on pelagic algal patterns and processes, neglecting the potential importance of benthic primary producers. While the role of benthic algae in lake primary production has recently gained recognition [28–34], conceptual and empirical studies of benthic algae have primarily focused on the roles of nutrients, water transparency, and hypsography [35, 36]. To our knowledge, the potential influence of the depth of the mixed surface layer has not been addressed in relation to benthic algae. Yet, it seems plausible that mixed layer depth should affect benthic algae through several mechanisms, e.g. both directly through its influence on the position of the thermocline (a breakpoint in temperature and concentrations of nutrients and CO_2_) in relation to the depth distribution of photic and aphotic lake bottoms, and indirectly through its effect on pelagic algae and, thus, the competitive impact of pelagic algae through shading. Understanding the effects of stratification and the depth of the mixed surface layer on the dynamics of benthic and pelagic algae therefore represents a significant gap of knowledge that we address in this paper using a conceptual modelling approach.

Given that benthic primary production in a lake strongly depends on the depth distribution of the bottom area above the photosynthetic compensation point [3, 33, 35], the exploration of the impact of mixed surface layer depth on the interaction between benthic and pelagic algae is only meaningful in the context of lake hypsography. Building on previous work [37], we therefore use a dynamical modeling approach — the two-dimensional, spatially explicit algal model ‘Lake2D’ — that faithfully represents the continuous depth distribution of the lake bottom and the water column above. Recently, we used Lake2D to explore algal dynamics under the simplifying assumption of vertically homogeneous turbulent mixing rates, intentionally neglecting stratification [38]. That study suggests that the strength of vertical mixing can have substantial impacts on benthic algae by mediating the competitive interactions between pelagic and benthic algae. To assess the potential impact of mixed surface layer depth on the dynamics of benthic and pelagic algae, we here apply Lake2D to lakes where the water is vertically stratified into two mixing regimes: a shallow regime with high turbulent mixing, and a deep regime with slow turbulent mixing, separated by a sharp boundary.

We then explore the impact of the depth of this boundary on pelagic and benthic algal dynamics across a comprehensive range of lake sizes, lake depths, and nutrient content. Our analysis provides new insights into the dynamics of lake primary production and the role of the mixed layer depth in structuring the spatial distribution of pelagic and benthic algae and the composition of total algal biomass across various lake types.

## 2 Model description

In the following section, we give a conceptual overview of the model ‘Lake2D’ as used in the present study. A schematic of Lake2D is presented in Fig. 1, with colored arrows illustrating the physical and biological processes included in the model. A detailed mathematical description of the model and parameter values is provided in Appendix A. Lake2D uses four partial differential equations and one algebraic equation to describe the rates of change of the following five spatially and temporally varying state variables: The concentrations of pelagic algal biomass (*A*) and a potentially growth-limiting inorganic nutrient (*N*_d_) in the water, the intensity of (potentially growth-limiting) sunlight (*I*) in the water, and the areal densities of sedimented particulate nutrients (*N*_s_) at the lake bottom and of benthic algal biomass (*B*) at the sediment-water interface. Algal biomass is expressed in units of carbon, and the nutrients in units of phosphorus (P). For simplicity, we assume that lakes are closed systems, with no external input or output of phosphorus. The total nutrient content of a lake is thus quantified by the temporally constant, lake-averaged total P per unit surface area (*N*_area_).

**Figure 1.**
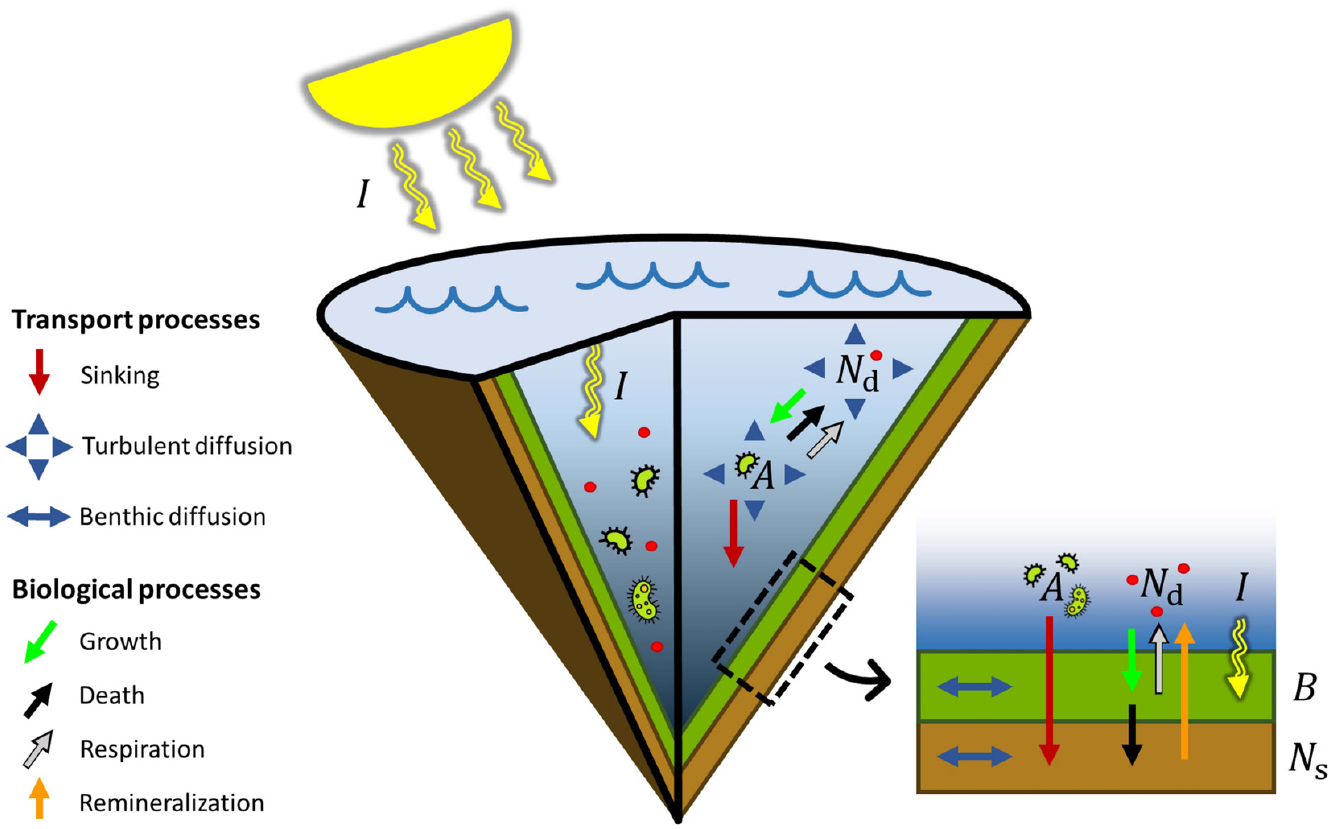
Schematic cross-section of the 2D algal model Lake2D for a cone-shaped lake. The model describes the dynamics of pelagic algae (A) and dissolved nutrients (N_d_) in the water, and the dynamics of benthic algae (B) on top of a layer of sedimented particulate nutrients (N_s_) on the bottom. In the water, Light (I) is attenuated by algae and non-algal background attenuation in the water, by benthic algae inside the benthic algal layer. Processes included in the model are indicated as follows: Red arrows represent sinking pelagic algae, and blue triangles represent turbulent mixing of pelagic algae and dissolved nutrients. Green arrows represent algae consuming dissolved nutrients. Benthic algae and sedimented nutrients diffuse slowly along the bottom, represented by blue double-arrows. Black and gray arrows represent algal nutrient losses from death and respiration, respectively. Finally, remineralization of nutrients from the sediment into the water is represented with an orange arrow.

A distinguishing feature of Lake2D is that it assumes radial symmetry of the lake and no directional bias in horizontal mixing. This assumption eliminates gradients in the azimuthal direction around the lake’s central axis, meaning that the values of all state variables are constant as one traverses the lake in a circle at a constant radial distance *r* from the lake center and a constant vertical depth *z* from the lake surface. The assumption thus effectively reduces the model’s spatial degrees of freedom from three to two, such that spatial dynamics can be fully represented with a 2-dimensional description of transport processes in the vertical (*z*) and radial (*r*) directions. While Lake2D can handle any radially symmetric basin shape, we assume a cone-shaped lake geometry with a linearly sloping bottom, because this simple topography facilitates visualization and interpretation of results, but is also representative of the hypsography of many real lakes [39–41].

The water column is vertically stratified into a well-mixed surface layer of thickness *z*_mix_ and a weakly mixed deeper layer below. Throughout the paper, we use the symbol 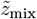 to express the depth of the thermocline in relative terms as a fraction of maximum lake depth (*z*_max_)

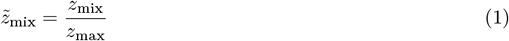

such that 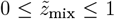. We simplify the thermocline to a sharp boundary, with high turbulent mixing above the thermocline and low turbulent mixing below. Turbulent mixing of pelagic algae and dissolved P is described as a diffusion process. Horizontal and vertical diffusion coefficients above and below the thermocline were set to constant values from the literature (Appendix A, Table 1). In addition to being turbulently mixed, pelagic algae sink with a constant velocity. When reaching the bottom, pelagic algae die and their nutrients are transferred to the pool of sedimented particulate nutrients, assuming a fixed phosphorus quota per unit biomass. Vertical light attenuation is modeled using Beer-Lambert’s Law. In the water column, light is attenuated by pelagic algae with a fixed, biomass-specific attenuation coefficient and by background turbidity representing light attenuation by non-algal materials and water itself.

**Table 1.**
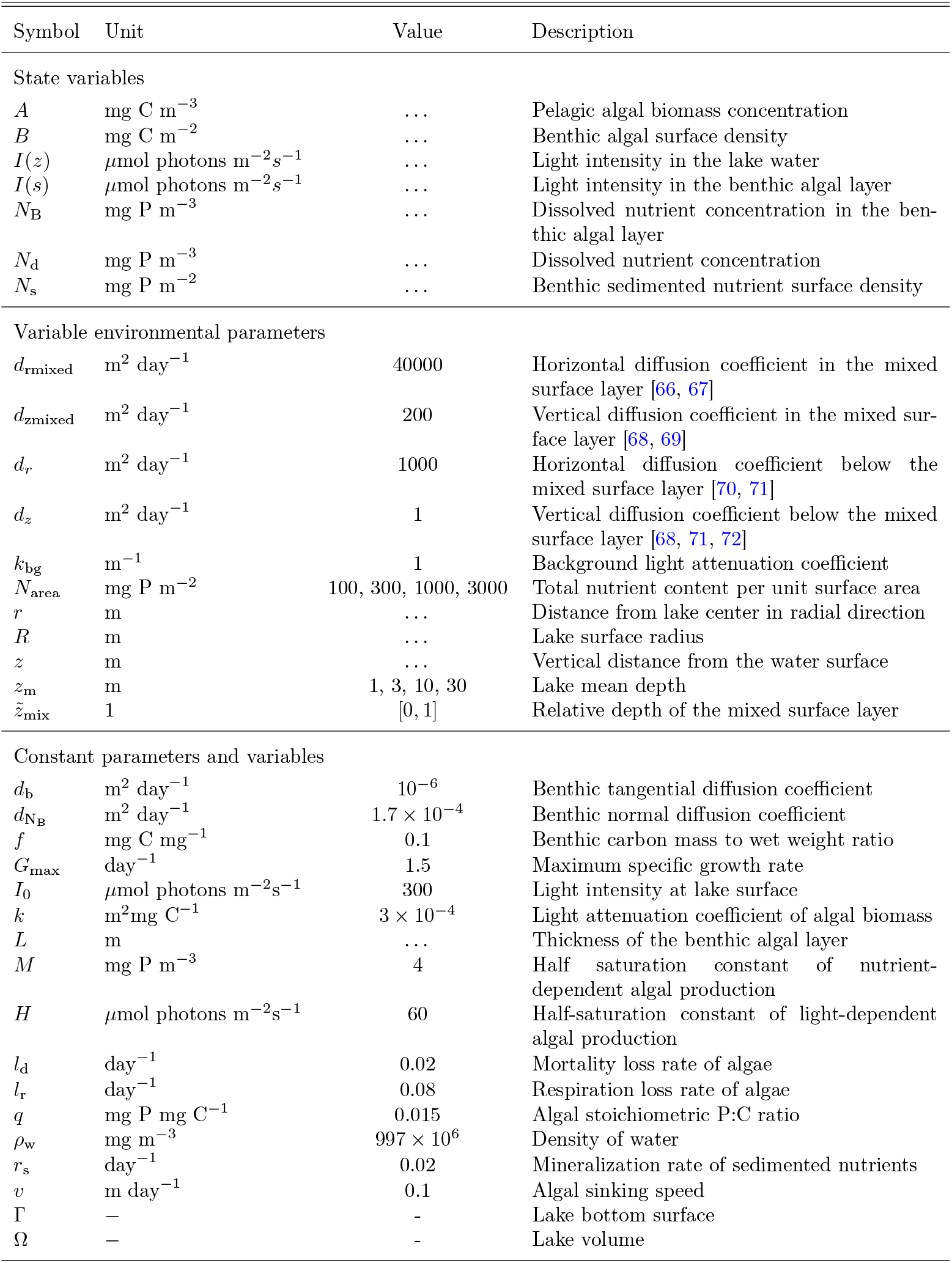
State variables and parameters. Note that the value chosen for the vertical diffusion coefficient in the mixed layer (d_rmixed_ = 200 m^2^ day^−1^), was intentionally set to a higher value than the higher end of the published range (≈ 35 m^2^ day^−1^). This deliberate choice was made to more clearly demonstrate the negative effect of a deep mixed layer on total pelagic biomass in deep, nutrient-poor lakes (Fig. 2, z_m_ = 30 m, N_area_ ≤ 300 mg P m^−2^). This higher value did not have any significant effect on the model’s dynamics in other regions of the explored parameter space.

At any point in the lake, algae take up nutrients in proportion to their growth rate, which is limited by either the local light intensity or the local concentration of dissolved P. This limitation is modeled using a minimum function, where light- and nutrient-limited growth are each represented by Monod functions. While the local concentration of dissolved P is modeled explicitly in the pelagic habitat, this is not the case in the benthic habitat. For each point along the lake bottom, we instead approximate the local vertical profile of dissolved P concentration inside the benthic algal layer using a separate differential equation (see Appendix A.2, eq. 15). We then use this concentration profile to determine the minimum of the light- and P-limited growth rate and the corresponding diffusive nutrient supply rate into the benthic algal layer, which we assume to come exclusively from the water immediately above. While nutrient uptake from the sediment is thus not explicitly considered, we make the simplifying assumption that sediment nutrients that have been mineralized (described with a first-order process) are released into the water directly above the benthic algal layer.

To focus our analyses on the impact of environmental factors, we assume that pelagic and benthic algae have identical biological parameters with respect to growth, light attenuation, and elemental stoichiometry, and that (density-independent) loss rates from respiration and mortality are the same for both algal types. Because algal carbon-to-phosphorus stoichiometry is fixed, carbon losses from respiration lead to corresponding losses of dissolved phosphorus into the water. Phosphorus from dead pelagic algae is instantaneously released into the water column in dissolved form, while phosphorus from dead benthic algae enters the pool of sedimented particulate nutrients. A minimal diffusive transport of benthic algae and sedimented nutrients in the radial direction along the bottom is included.

## 3 Methods

### Design of numerical experiments

We conducted extensive numerical simulations of Lake2D to evaluate the influence of mixed surface layer depth on algal dynamics across a comprehensive range of environmental conditions. Four environmental parameters were systematically varied: Lake mean depth (*z*_m_), total nutrient content per surface area (*N*_area_), lake surface area, and relative thermocline depth 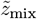. Lake area and 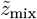 constitute a two-dimensional slice of the parameter space, which we will refer to as the lake area-mixed layer space. Similarly, the parameters *N*_area_ and *z*_m_ form a separate two-dimensional slice of the parameter space, which we refer to as the nutrient-depth space. Collectively, *z*_m_, *N*_area_, Lake area, and 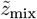 define a 4-dimensional environmental parameter space, which we numerically explored in a fully factorial design, varying the parameters over ranges that are representative of the vast majority of the world’s lakes [42–45]. Specifically, we performed 21904 numerical simulations resulting from the combination of four values of *z*_m_ (1, 3, 10, and 30 m), four values of *N*_area_ (100, 300, 1000, and 3000 mg P m^*−*2^), and a 37 *×* 37 factorial lake area-mixed layer space, where lake area was varied on a logarithmic scale over the interval [0.01, 1000] km^2^ (representing 99.99% of the world’s lakes *>* 0.01 km^2^ [46]), and 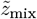 was varied linearly on the interval [0, 1] (covering the full range of potential mixed layer depths).

### Execution of numerical experiments

Lake2D was implemented in MATLAB using a finite volume method, employing a diagonally cut grid with a minimum resolution of 30×30 equally spaced grid cells. The model was integrated in time using MATLAB’s built-in ODE15s function. To maintain computational accuracy, the local grid resolution was increased as necessary near steep spatial gradients, which can arise at specific depths or distances from the lake center under certain environmental conditions. All simulations were run to steady state from the same initial condition: 98% of total nutrients in dissolved form, 1% in pelagic algae, and the remaining 1% in benthic algae, all evenly distributed across the lake’s volume and bottom, respectively. With few exceptions, simulations converged to within less than *±* 5% of steady state conditions within < 100 days.

### Model analyses and output visualization

We began our analyses of model output at the whole-lake scale, with an emphasis on the relationship between the relative thermocline depth 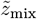 and algal dynamics. In a first step, we integrated steady-state pelagic, benthic, and total (pelagic + benthic) algal biomass over the entire lake, expressed as average biomass per unit lake surface area. To visualize these results in the explored 4-dimensional parameter space, we produced for each combination of *N*_area_ and *z*_m_ a heat map in lake area-mixed layer space of the steady-state total (*A* + *B*), pelagic (*A*), and benthic (*B*) algal biomass in lake area-mixed layer space (Fig. 2).

**Figure 2.**
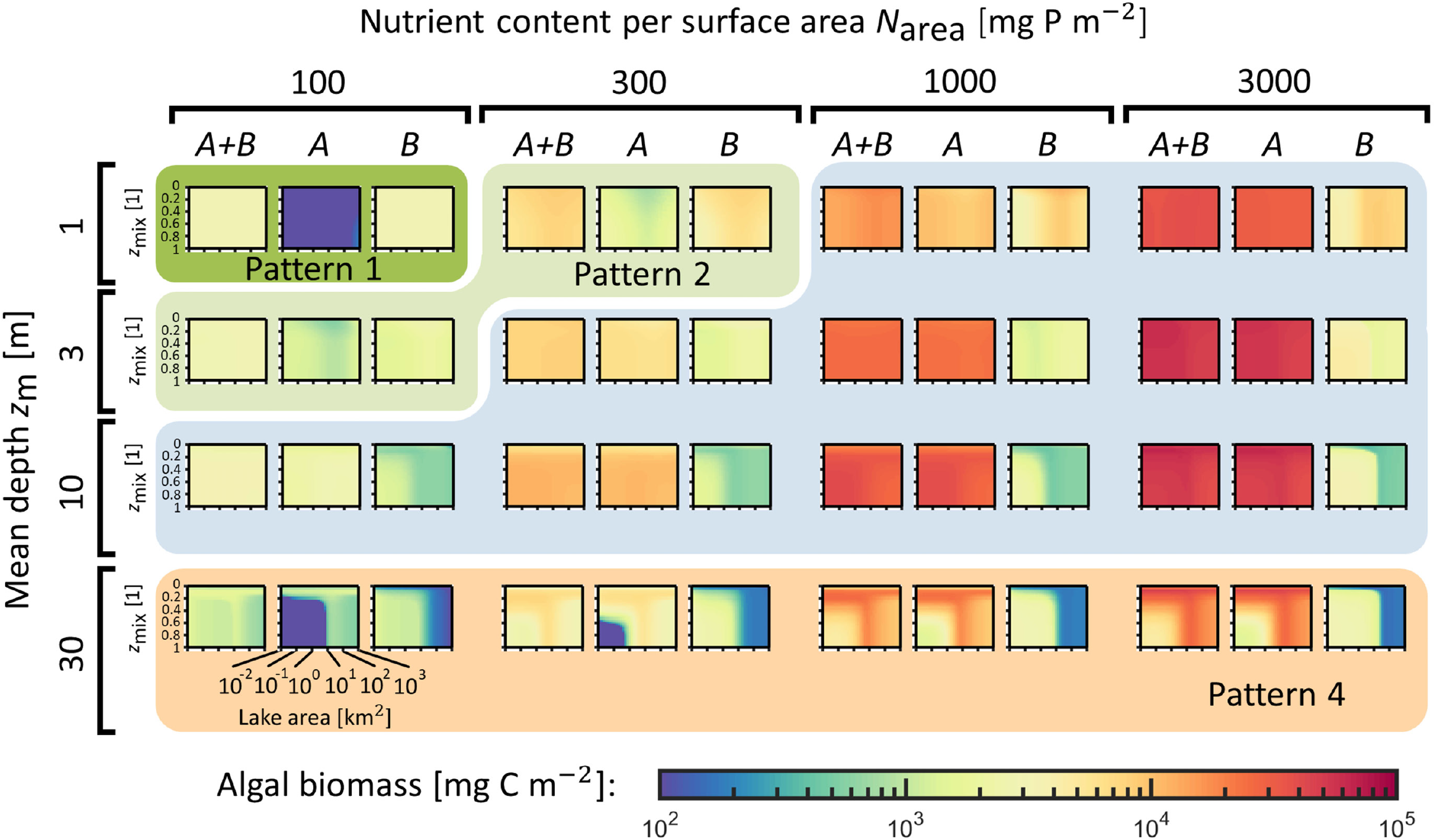
(Previous page.) Heat maps of lake-wide pelagic (A), benthic (B), and total (A + B) algal biomass for varying total nutrient content per surface area N_area_, lake mean depth z_m_, lake area, and relative mixing depth 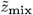. Lake area is varied along the horizontal axis of each panel in the range [10^−2^, 10^3^] km^2^, and the relative mixing depth 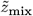 is varied along the vertical axis of each panel in the range [0, 1]. The vertical and horizontal axes are plotted on a linear scale and log10 scale respectively. Each pixel in a panel is the average biomass per unit lake surface area (mg C m^−2^) per unit lake surface area in log scale, see the colorbar at the bottom of the figure. Panels with a dark green background correspond to Pattern 1 simulations; panels with a light green background correspond to Pattern 2 simulations; panels with a blue background correspond to Pattern 3 simulations; and panels with an orange background correspond to Pattern 4 simulations.

Next, we analyzed the influence of the four environmental drivers on the spatial distribution of algae within lakes, and how these dynamics in turn give rise to the observed patterns at the whole-lake scale. To this end, we present individual simulations of two lake sizes (0.3 km^2^ and 300 km^2^) and two relative thermocline depths 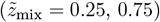, across different values of nutrient content *N*_area_ and mean depth *z*_m_ (Fig. 3-6). We finally examined the influence of the four environmental drivers on the partitioning of a lake’s total nutrients between benthic and pelagic biomass and phosphorus pools (Fig. 7).

**Figure 3.**
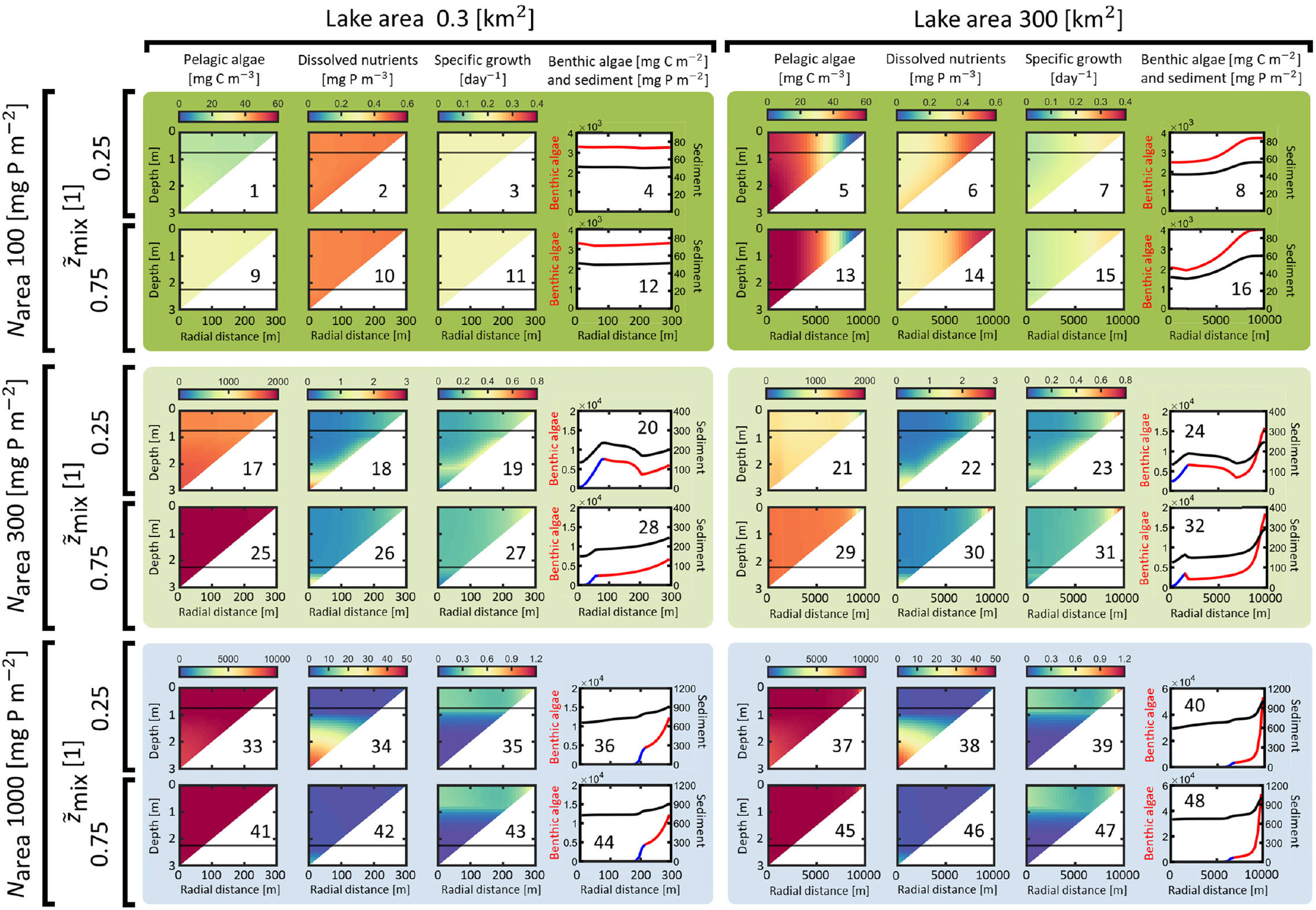
Twelve example simulations visualizing Lake2D model output for lake mean depth z_m_ = 1 m and a factorial combination of two relative mixed layer depths (0.25 and 0.75, inner brackets on the left) nested inside three nutrient levels (100, 300, 1000 mg P m^2^, outer brackets on the left), and two lake sizes (0.3 and 300 km^2^, brackets on top). Each set of four horizontally grouped panels (e.g., panels 1-4, 5-8, 9-12, etc.) displays the output of a single Lake2D simulation., displaying the steady state spatial distributions of (from left to right): the concentration of pelagic algal biomass, the concentration of dissolved phosphorus, the specific growth of pelagic algae, and the surface density of benthic algal biomass and sedimented phosphorus, respectively (see the headings above each column of panels). Horizontal lines in triangular plots indicate the position of the lower boundary of the mixed layer. Background colors are coded as in Fig. 2, where dark green indicates Pattern 1, light green indicates Pattern 2, and blue indicates Pattern 3.

## 4 Results

Whole-lake pelagic, benthic, and total algal biomass across the explored 4-dimensional parameter space is presented in Fig. 2. First, we analyze algal dynamics on this whole-lake scale, and discern the influence of the thermocline depth 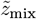 on whole-lake algal dynamics. We then analyze how 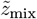 affects the spatial distribution of algal biomass within individual lakes (Fig. 3-6), and explain when and how these spatial dynamics give rise to the patterns in whole-lake biomass. Finally, we present plots of the contribution of pelagic algae, benthic algae, dissolved nutrients, and sedimented nutrients to the total nutrient pool, and examine the impact of 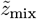 on the composition of total nutrients for varying 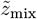 across a range of lake sizes and total nutrient content (Fig. 7).

### 4.1 Whole-lake algal biomass dynamics

#### Pattern 1

In the shallow, nutrient-poor Pattern 1 lakes (Fig. 2, *N*_area_ = 100 and *z*_m_ = 1. Panels on a dark-green background), significant nutrient-limitation restricts the growth of pelagic algae. This low growth, combined with sinking losses, prevents the algae from accumulating, leading to an overall low concentration of pelagic algal biomass throughout (dark blue pelagic algal biomass panel *A*). In contrast, benthic algae thrive in these shallow systems and constitute the majority of total biomass (compare the benthic algal biomass panel *B* to the combined biomass panel *A* + *B*). In Pattern 1 lakes, neither 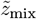 nor lake size has any effect on whole-lake algal biomass in lake area-mixed layer space.

#### Pattern 2

Pattern 2 occurs in slightly deeper or more nutrient-rich lakes (Fig. 2, *N*_area_ = 100, *z*_m_ = 3 and *N*_area_ = 300, *z*_m_ = 1. Panels on a light-green background). In these lakes, pelagic algal biomass is significantly higher compared to Pattern 1 lakes. This is due either to reduced sinking losses from increased depth (*z*_m_ = 3), or increased growth from higher nutrient content (*N*_area_ = 300), both of which allow pelagic algal growth to compensate for sinking losses. Akin to Pattern 1, whole-lake dynamics in Pattern 2 lakes are characterized by little variation in lake area-mixed layer space, with benthic algae constituting the majority of total biomass.

#### Pattern 3

Increasing the nutrient content or the mean depth further results in another qualitative shift in whole-lake dynamics to a new regime in nutrient-depth space, which we call Pattern 3 (Fig. 2, *z*_m_ *≤* 10, panels on a blue background). In these lakes, pelagic algal biomass is higher compared to Pattern 1 and Pattern 2 lakes, and now constitutes the majority of total biomass. Moreover, pelagic algal biomass increases with increasing *N*_area_, due to the increased growth of nutrient-limited algae in well-lit shallow areas (compare Pattern 2 pelagic algal panels *A* for increasing *N*_area_ in Fig 2). Conversely, benthic algal biomass does not show the same positive correlation with the total nutrient content in these systems. Furthermore, whole-lake pelagic algal biomass shows little to no change or a slight increase with increasing *z*_m_, whereas whole-lake benthic algal biomass decreases as *z*_m_ increases due to overall poorer light conditions. Akin to Pattern 1 and 2 lakes, whole-lake biomass in Pattern 3 systems generally does not vary much in lake area-mixed layer space, with one notable exception: for *z*_m_ = 10 m, whole-lake benthic algal biomass displays a distinct L-shaped Pattern in lake area-mixed layer space, with highest biomass occurring for lake areas *≤* 10 km^2^ and 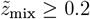. Note that the size of this high biomass area in lake area-mixed layer space increases with *N*_area_, while the value of whole-lake benthic biomass inside and outside this region is independent of *N*_area_.

#### Pattern 4

In deep lakes (Fig. 2, *z*_m_ = 30, panels on an orange background), algal dynamics exhibit a fourth unique pattern, which we call Pattern 4. In this regime, pelagic and benthic algal biomass display a characteristic L-shaped pattern in lake area-mixed layer space, where whole-lake pelagic algal biomass is notably lower under low-area, high-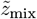 conditions compared to the rest of lake area-mixed layer space. This reduction is due to the rapid transport of algae from well-lit, shallow areas to aphotic depths. As *N*_area_ increases, increased pelagic algal growth compensates for these losses, resulting in higher pelagic algal biomass in low-area, high-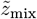 conditions (compare bottom left corner of Pattern 3 *A* panels in Fig. 2 for *N*_area_ *≤* 300 and *N*_area_ *≥*1000). Conversely, benthic algal biomass displays the opposite pattern to pelagic algal biomass, with higher biomass concentrations for low-area, high-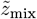 conditions compared to the remainder of lake area-mixed layer space, akin to the patterns in benthic algal biomass in *z*_m_ = 10 Pattern 2 lakes.

The complementary L-shaped patterns for pelagic and benthic algal biomass in these systems give rise to distinct whole-lake biomass compositions. For lakes with *N*_area_ *≤* 300, benthic algae account for the majority of the total biomass under low-area, high-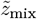 conditions. However, in the rest of lake area-mixed layer space, pelagic algae become the largest contributor to the total biomass. When *N*_area_ *≥* 1000, pelagic algal biomass is high enough even in unfavorable low-area, high-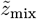 conditions, leading to pelagic algae contributing most to the total biomass across the entire lake area-mixed layer space. To fully understand the cause of these L-shaped patterns requires the analysis of the spatial dynamics of algae across lake area-mixed layer space for these systems.

### 4.2 Spatial dynamics within lakes

#### Pattern 1

In the small Pattern 1 lakes (Fig. 3, panel 1-4, 9-12), the lake water is completely well mixed, regardless of thermocline depth 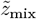. As a result, these lakes exhibit no discernible horizontal or vertical gradients in the concentrations of pelagic algae or dissolved nutrients (Fig. 3, uniform color in panel 1-3 and 9-11). Algal growth is completely nutrient-limited due to the low total nutrient content and favorable light conditions, and together with the well-mixed dissolved nutrients results in a uniform areal density of benthic algal biomass (Fig. 3, panel 4, 12).

In contrast, large Pattern 1 lakes (Fig. 3, panel 5-8, 13-16) exhibit horizontal gradients in pelagic algae and dissolved nutrients, enabled by slow horizontal turbulent mixing. Here, pelagic algal concentration is lower in areas close to the lake shore, and higher in the middle of the lake (Fig. 3, panel 5, 13). Dissolved nutrients display the opposite pattern, exhibiting the highest concentrations in the middle of the lake (Fig. 3, panel 6, 14). This is caused by high pelagic algal sinking losses in shallow areas, which the relatively slow horizontal transport of pelagic algae from deeper areas and local growth cannot fully compensate for. Consequently, pelagic algal concentrations are low in nearshore areas, even though the specific growth rate of pelagic algae is highest there (Fig. 3, panel 7, 15). Akin to the small Pattern 1 lakes, in large Pattern 1 lakes algal growth is completely nutrient limited. However, benthic algae now exhibit the highest areal densities along the lake shore where the concentrations of dissolved nutrients are highest (Fig. 3, panel 8, 16). Furthermore, 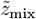 has no effect on spatial dynamics in large Pattern 1 lakes, due to the lack of vertical pelagic gradients regardless of thermocline depth.

#### Pattern 2

In the slightly deeper or more nutrient-rich Pattern 2 lakes, shading from pelagic algal biomass is higher compared to Pattern 1 lakes, with light-limited algal growth in deep areas and nutrient-limited growth in shallow areas. The vertical gradient in pelagic algal specific growth and the change in color in the benthic algal line plots reflect this spatial pattern (Fig. 3, 4).

**Figure 4.**
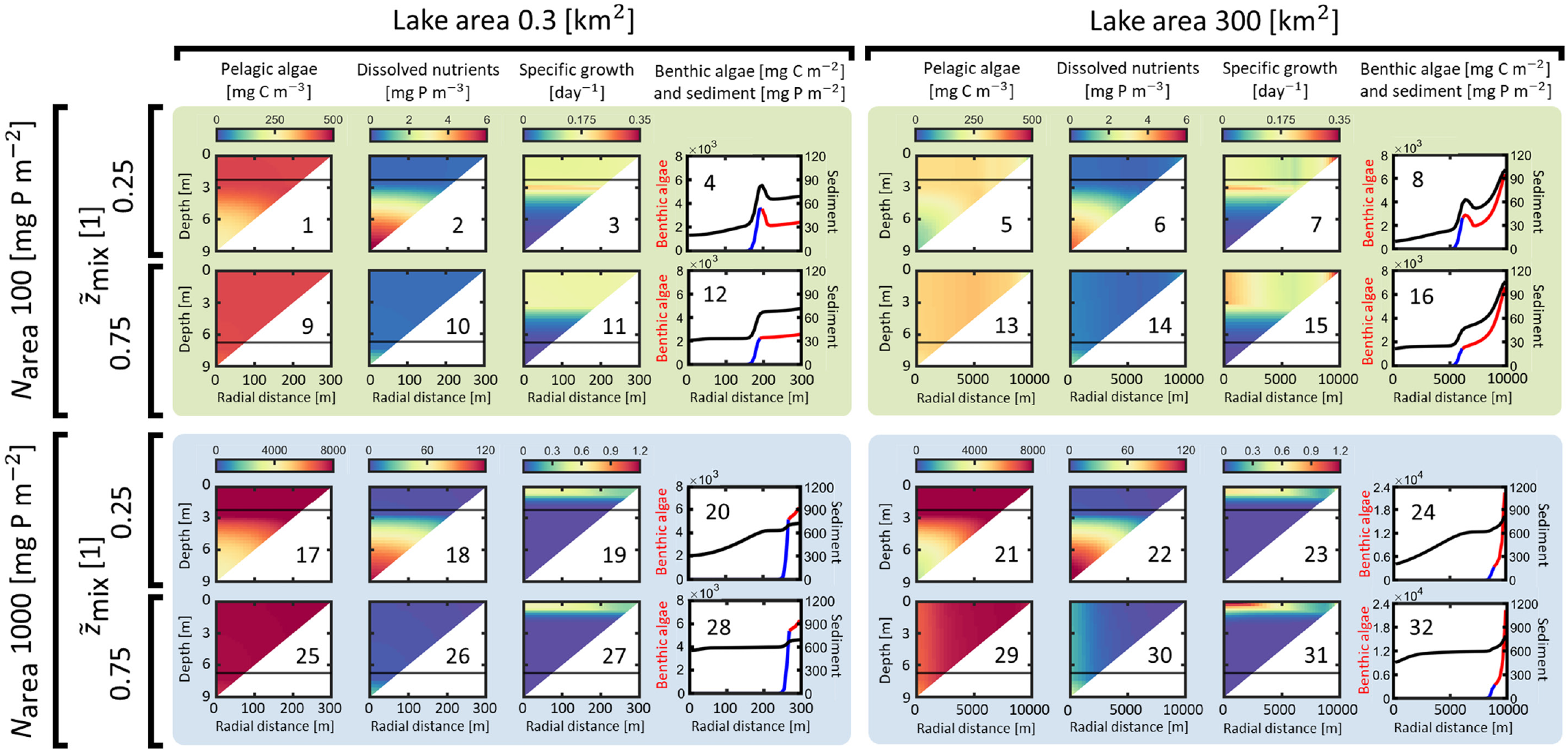
Eight example simulations visualizing Lake2D model output for lake mean depth z_m_ = 3 m and a factorial combination of two relative mixed layer depths (0.25 and 0.75, inner brackets on the left) nested inside three nutrient levels (100, 300, 1000 mg P m^2^, outer brackets on the left), and two lake sizes (0.3 and 300 km^2^, brackets on top). Each set of four horizontally grouped panels (e.g., panels 1-4, 5-8, 9-12, etc.) displays the output of a single Lake2D simulation., displaying the steady state spatial distributions of (from left to right): the concentration of pelagic algal biomass, the concentration of dissolved phosphorus, the specific growth of pelagic algae, and the surface density of benthic algal biomass and sedimented phosphorus, respectively (see the headings above each column of panels). Horizontal lines in triangular plots indicate the position of the lower boundary of the mixed layer. The background color is coded as in Fig. 2, where light green indicates Pattern 2, and blue indicates Pattern 3.

In shallow Pattern 2 lakes, there are no significant gradients in the concentration of pelagic algae (Fig. 3, panel 17, 21, 25, 29), and 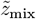 has little effect on the spatial dynamics of pelagic algae. Dissolved nutrients accumulate at the bottom of the lake, with a more pronounced vertical gradient when the thermocline is shallow (Fig. 3, panel 18, 26).

In deep Pattern 2 lakes, a change in thermocline depth introduces qualitative differences in the spatial dynamics of the lake. When the thermocline is deep, the water is well mixed, and no significant vertical gradients of pelagic algae or dissolved nutrients are present (Fig. 4, panel 9-10, 13-14). Conversely, when the thermocline is shallow, distinct vertical gradients form, with lower pelagic algal concentrations at the bottom and a corresponding accumulation of dissolved nutrients in the deeper water (Fig. 4, panel 1-2, 5-6). Consequently, 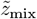 influences the spatial dynamics in these systems by controlling the formation of these vertical gradients.

In small Pattern 2 lakes with shallow thermoclines, benthic algal biomass exhibits a unique phenomenon: a maximum in benthic algal biomass occurs at an intermediate depth. This peak corresponds to the transition from light to nutrient limitation (Fig. 3, panel 20, 24, Fig. 4, panel 4, 8), and requires opposing vertical gradients of light and dissolved nutrients in the water, such that algal specific growth displays a local (and sometimes global) maximum at an intermediate depth. Moreover, if the thermocline is too deep in small lakes, the resulting fast turbulent mixing combined with the small lake size results in negligible gradients in dissolved nutrients at depths where benthic algae are nutrient-limited (Fig. 3, panel 26, and Fig. 4, panel 14). Consequently, no benthic deep chlorophyll maximum is observed (Fig. 3, panel 28, and Fig. 4, panel 16).

#### Pattern 3

In shallow Pattern 3 lakes, pelagic algae display no discernible spatial gradients, regardless of thermocline depth 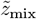 (Fig. 3, panel 33, 37, 41, 45). Dissolved nutrients, however, do show a dependence on 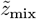: they are well mixed when the thermocline is deep, but accumulate in deeper areas when the thermocline is shallow (Fig. 3, panel 34, 38, 42, 46). Lake size has no discernible effect on the distribution of either pelagic algae or dissolved nutrients in these shallow Pattern 3 systems.

In moderate-depth Pattern 3 lakes, the patterns for dissolved nutrients remain identical to the shallower lakes, with bottom accumulation for shallow thermoclines and well-mixed conditions for deep thermoclines (Fig. 4, panel 18, 22, 26, 30). Similarly, lake size does not affect their spatial distribution. Pelagic algae, however, now exhibit a vertical gradient for shallow thermoclines, with concentrations highest at the surface, while remaining well mixed when the thermocline is deep (Fig. 4, panel 17, 21, 25, 29).

A more complex spatial dynamic emerges in deep Pattern 3 lakes, where lake size significantly influences spatial dynamics. In small lakes, both pelagic algae and dissolved nutrients are horizontally well mixed (Fig. 5, panel 1, 2, 9, 10, 17, 18, 25, 26). For shallow thermoclines, a vertical gradient forms, with pelagic algae concentrated near the surface and dissolved nutrients accumulating in deeper areas (Fig. 5, panel 1, 2, 17, 18). As the thermocline deepens, these gradients disappear (Fig. 5, panel 9, 10, 25, 26).

**Figure 5.**
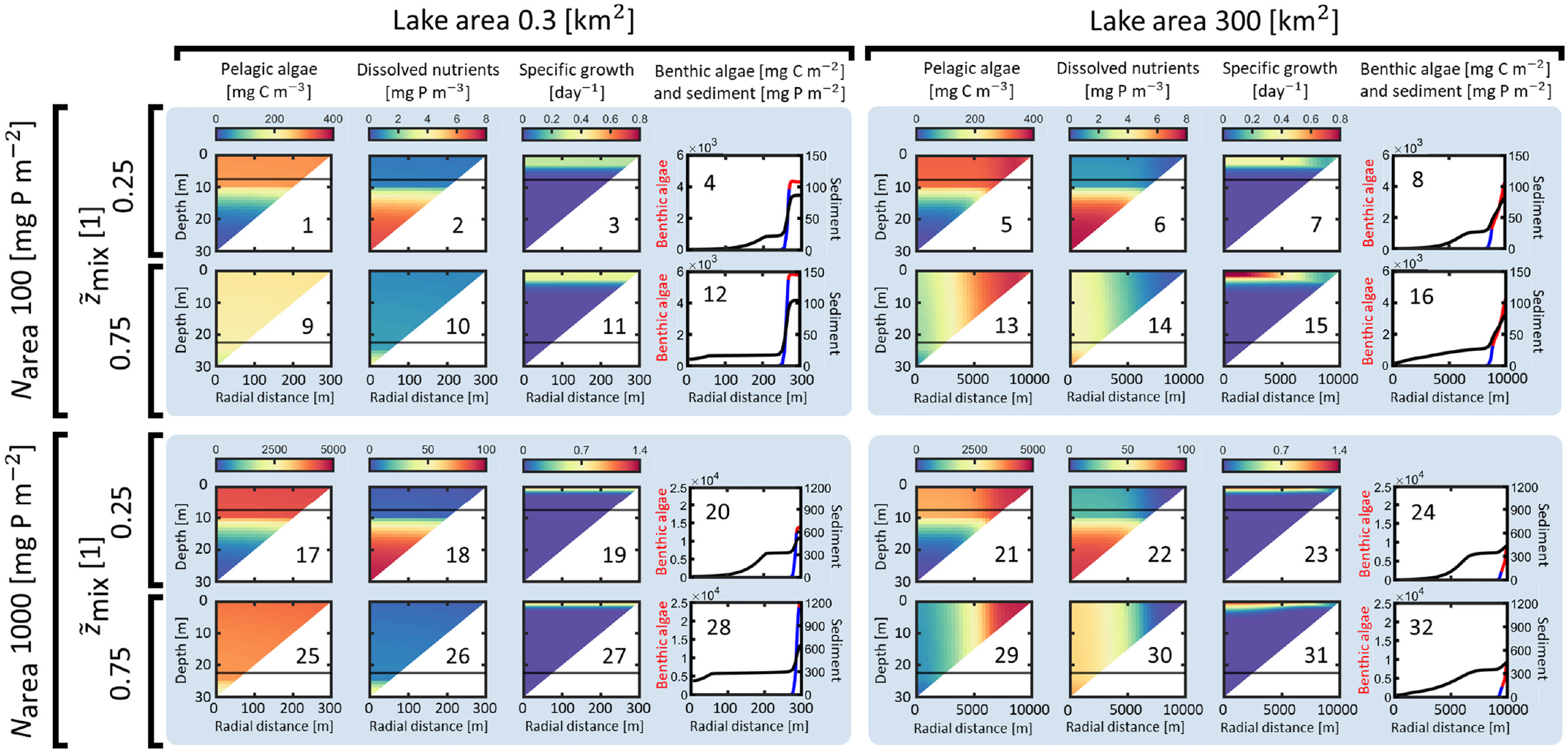
Eight example simulations visualizing Lake2D model output for lake mean depth z_m_ = 10 m and a factorial combination of two relative mixed layer depths (0.25 and 0.75, inner brackets on the left) nested inside three nutrient levels (100, 300, 1000 mg P m^2^, outer brackets on the left), and two lake sizes (0.3 and 300 km^2^, brackets on top). Each set of four horizontally grouped panels (e.g., panels 1-4, 5-8, 9-12, etc.) displays the output of a single Lake2D simulation., displaying the steady state spatial distributions of (from left to right): the concentration of pelagic algal biomass, the concentration of dissolved phosphorus, the specific growth of pelagic algae, and the surface density of benthic algal biomass and sedimented phosphorus, respectively (see the headings above each column of panels). Horizontal lines in triangular plots indicate the position of the lower boundary of the mixed layer. Background colors are coded as in Fig. 2, where a blue background color indicates Pattern 3.

The spatial dynamics in larger lakes differ significantly with 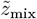. With a shallow thermocline, large Pattern 3 lakes mirror small Pattern 3 lakes, with negligible horizontal gradients and vertical stratification of pelagic algae and dissolved nutrients (Fig. 5, panel 5 6, 21, 22). However, when the thermocline is deep, a distinct horizontal gradient develops. Pelagic algal concentrations are highest along the lake shore and lowest in the middle, while dissolved nutrients display the opposite pattern (Fig. 5, panel 13, 14, 29, 30). In all Pattern 3 lakes, benthic algae are restricted to nearshore areas due to light limitation in deeper regions. Their distribution show no qualitative change with variations in lake size or thermocline depth.

#### Pattern 4

The conditions in the deep Pattern 4 lakes are characterized by poor light conditions, with pelagic algal growth relegated to areas close to the lake surface (Fig. 6, specific growth panels). Likewise, Benthic algae only grow in the shallowest areas of lakes with the highest areal densities found right along the lake shore, analogous to Pattern 2 systems (Fig. 6, benthic algal panels). When the nutrient content is low and the lake size is small, the limited pelagic algal growth in Pattern 4 lakes cannot compensate for losses at aphotic depths, resulting in the extinction of pelagic algae (Fig. 6, panel 1, 9), where survival of pelagic algae only occurs for very shallow thermoclines (not shown).

**Figure 6.**
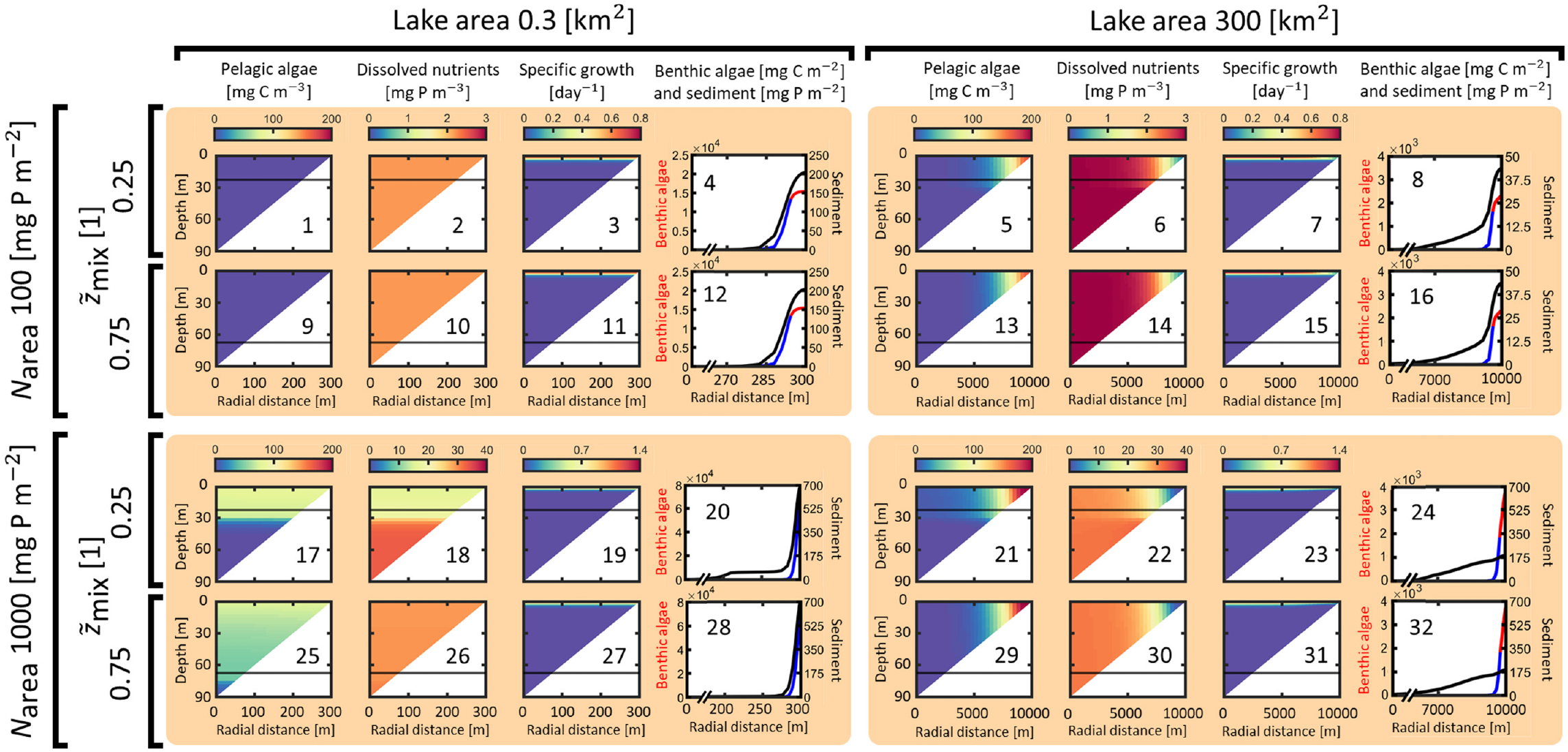
Eight example simulations visualizing Lake2D model output for lake mean depth z_m_ = 30 m and a factorial combination of two relative mixed layer depths (0.25 and 0.75, inner brackets on the left) nested inside three nutrient levels (100, 300, 1000 mg P m^2^, outer brackets on the left), and two lake sizes (0.3 and 300 km^2^, brackets on top). Each set of four horizontally grouped panels (e.g., panels 1-4, 5-8, 9-12, etc.) displays the output of a single Lake2D simulation., displaying the steady state spatial distributions of (from left to right): the concentration of pelagic algal biomass, the concentration of dissolved phosphorus, the specific growth of pelagic algae, and the surface density of benthic algal biomass and sedimented phosphorus, respectively (see the headings above each column of panels). Horizontal lines in triangular plots indicate the position of the lower boundary of the mixed layer. Background colors are coded as in Fig. 2, where an orange background color indicates Pattern 4.

**Figure 7.**
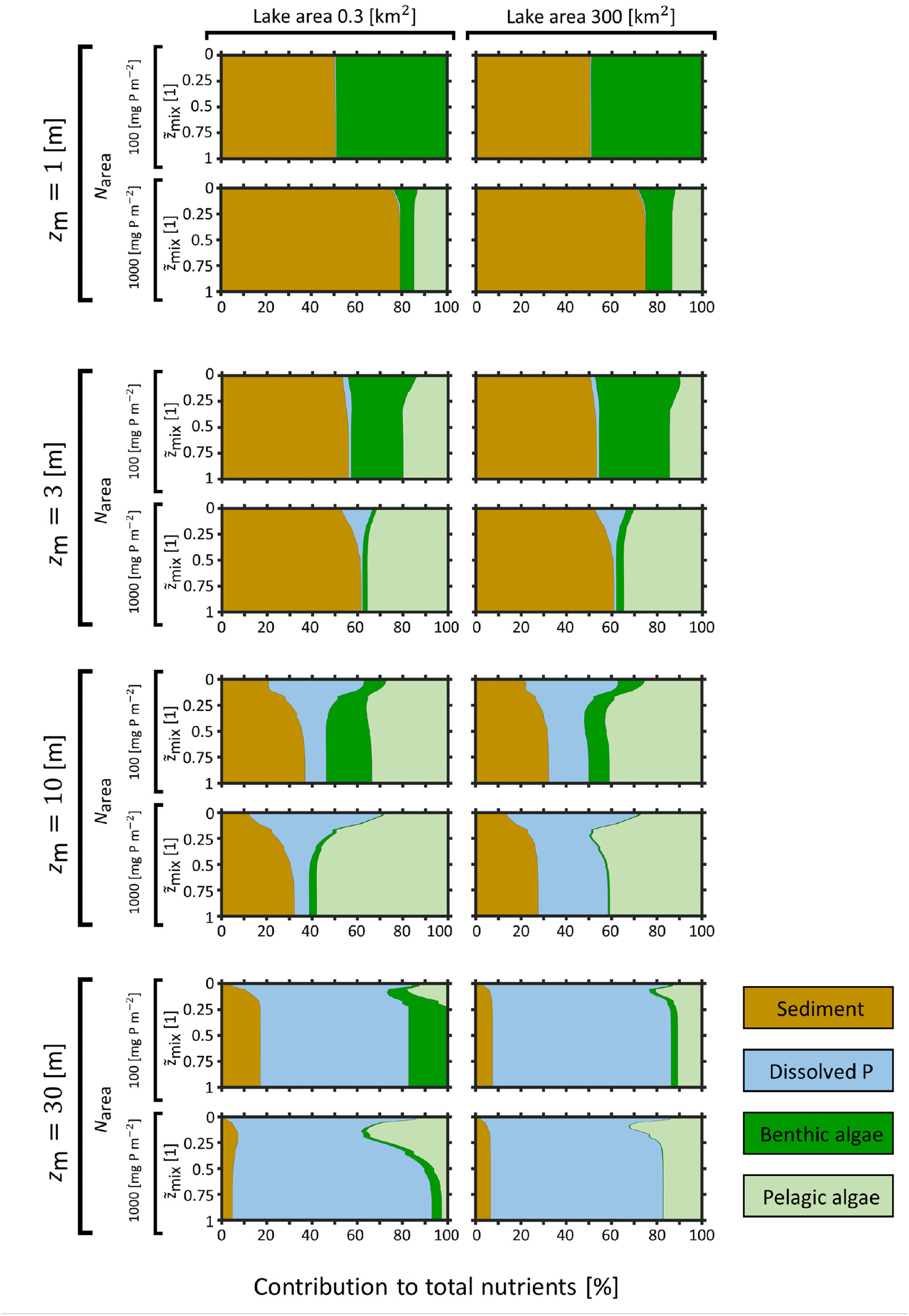
(Previous page.) Fractional contributions of pelagic algal biomass, benthic algal biomass, dissolved phosphorus, and sedimented phosphorus to a lake’s total phosphorus pool as a function of lake mean depth z_m_ (outer top brackets above each group of four panels), lake area (inner top brackets), nutrient content per surface area Narea (brackets on the left), and relative mixied layer depth 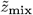 (y-axes). Each panel represents a vertical slice across the full range of relative mixed layer depths for lake sizes 0.3 and 300 km^2^, respectively, in the corresponding panels of Fig. 2.

In large Pattern 4 lakes, slow horizontal turbulent mixing allows for horizontal gradients to form in pelagic algae and dissolved nutrients above the thermocline (Fig. 6, panel 5-6, 13-14, 21-22, 29-30). Here, pelagic algae survive in near-shore areas where the lake is shallow enough such that despite being well mixed vertically, pelagic algal growth can compensate for sinking losses, sustaining a local population of pelagic algae. Dissolved nutrients display the opposite pattern to the pelagic algal biomass, with the highest concentrations found in the middle of the lake, decreasing horizontally towards the lake shore (Fig. 6, panel 6, 14, 22, 30).

Finally, in small Pattern 4 lakes with high nutrient content, pelagic algae sustain a population across the entire surface of the lake for small 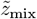 (Fig. 6, panel 17). Dissolved nutrients display a complementary trend with highest concentrations at aphotic depths, decreasing towards the surface (Fig. 6, panel 18). Here, 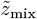 has a clear impact on spatial dynamics, resulting in pelagic algae being mixed downwards and dissolved nutrients being mixed upwards with increasing 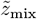 (Fig. 6, compare panel 17 with 25, and 18 with 26). Pelagic algal biomass decreases with increasing 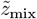, but unlike systems with less total nutrients, pelagic algae do not go extinct even for deep thermoclines (Fig. 6, panel 25).

Note that large, nutrient-rich Pattern 4 lakes with a shallow 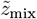 have higher dissolved nutrient concentrations in the middle of the lake compared to smaller, Pattern 4 lakes with the same nutrient content (Fig. 6, panel 17, 22). In spite of this, the larger lakes have much lower pelagic algal biomass in the middle of the lake compared to the smaller lakes (Fig. 6, panel 17, 21). This seeming contradiction reveals that the population of algae above deep areas in small, nutrient-rich Pattern 4 lakes is dependent on the influx of algae from shallow areas along the lake shore, which, due to the circular lake shape, are much larger compared to the middle of the lake.

Together, the survival of pelagic algae in shallow areas in large lakes and the along the lake surface in small lakes with shallow thermoclines, combined with a reduction (or extinction) of pelagic algal biomass in small lakes with deep thermoclines, gives rise to the L-shaped pattern observed in whole-lake algal biomass for Pattern 4 lakes (Fig. 2, *z*_m_ = 30 m).

### 4.3 Whole-lake composition of total nutrients

Fig. 7 illustrates the contribution of pelagic algal biomass, benthic algal biomass, dissolved nutrients, and sedimented nutrients to the total nutrient pool across varying 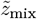, *N*_area_, and *z*_m_, which covers the full spectrum of observed trends across the four environmental drivers. Overall, *z*_m_ has the biggest impact on Whole-lake nutrient composition, followed by *N*_area_. In contrast, lake size has a smaller effect on the composition of the total nutrient pool, particularly in shallow lakes with *z*_m_ *≤* 3 m. In general, the effect of 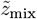 on the nutrient composition is the largest for 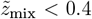, above which the nutrient composition stabilizes and does not change much with 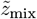.

The nutrient composition in lakes with *z*_m_ *≤* 3 m is characterized by a dominance of sedimented nutrients and a small contribution from dissolved nutrients. In these lakes, the increased survival of pelagic algae with increasing *N*_area_ results in a corresponding increase in the portion of sedimented nutrients due to increased sinking losses of pelagic algae, and a decrease in the portion of benthic algae. The thermocline depth 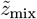 only affects nutrient composition for values of 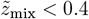.

In this range, the portion of benthic algae and dissolved nutrients is highest when 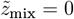. As 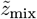 increases, these portions decrease, with a corresponding increase in the portion of sedimented nutrients and pelagic algae. For 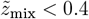, these portions do not change.

For lakes with *z*_m_ *≥* 10, the portion of total nutrients in the sediment and benthic algae is smaller compared to *z*_m_ *≤* 3 m lakes, whereas the portion of total nutrients in dissolved form is much higher. This is due to the accumulation of nutrients at aphotic depths in these deeper systems, where algae cannot grow due to light limitation. Moreover, the contribution of pelagic algae to total nutrients decreases with increasing *z*_m_, the opposite of the trend found in *z*_m_ *≤* 3 m lakes. Together, these two trends result in a unimodal pattern in the contribution of pelagic algae to the total nutrient pool for varying *z*_m_, peaking at *z*_m_ = 10 with lower contributions to the total nutrient pool in deeper and shallower lakes. Conversely, the contribution of benthic algae does not exhibit the same behavior, and decreases with increasing *z*_m_.

Similar to *z*_m_ *≤* 3 m lakes, in lakes with *z*_m_ *≥* 10 m 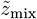 has the biggest effect on nutrient composition when 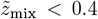, and the lakes exhibit an initial decrease in the portion of dissolved nutrients with increasing 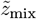 accompanied by a corresponding increase in the portion of sedimented nutrients and pelagic algae. However, for *z*_m_ = 30, the portion of total nutrients in pelagic algae does not stabilize around 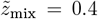. Instead, it increases at first for shallow 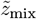, and then begins to decrease when 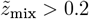. This is caused by the interplay of upwards turbulent mixing of dissolved nutrients increasing algal growth in shallow areas, and the downwards turbulent mixing of pelagic algae to aphotic depths increasing pelagic algal losses. For 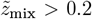, the pelagic algal losses from increased turbulent mixing outweigh the benefit of increased upwards transport of nutrients, resulting in a net decrease of pelagic algae.

## 5 Discussion

In this study, we used the two-dimensional, spatially explicit algal model Lake2D to comprehensively evaluate how a lake’s physical characteristics, specifically lake size (surface area) and the relative depth of the mixed surface layer 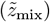, influence the ecological dynamics of both pelagic and benthic algae. Our findings both confirm, contextualize, and expand upon several well-established ecological principles, providing new insights into the dynamics of lake primary production. This study addresses a significant gap in our understanding of how the depth of the mixed surface layer and lake size influence the ecological dynamics of both pelagic and benthic algae across a wide range of environmental conditions.

### 5.1 Extending conceptual frameworks for pelagic algae

Our model results reproduce and contextualize established findings from classical one-dimensional models of pelagic algal dynamics while accounting for horizontal heterogeneity and varying lake depths. The classic critical depth principle [19], which states that phytoplankton blooms can only occur when the mixed layer is shallower than a critical depth, and the concept of critical turbulence, which states that phytoplankton blooms cannot not develop if the rate of turbulent mixing is above a critical level, are confirmed under certain conditions in our simulations: in small, deep Pattern 4 lakes with low nutrient content, a sufficiently deep mixed layer transports pelagic algae to aphotic depths, where losses cannot be compensated for by algal growth in shallow, sunlit surface waters, leading to their extinction (Fig. 2, *z*_m_ = 30, *N*_area_ *≤* 300). This aligns with the concept that a deeper mixed layer leads to greater light limitation and reduced algal growth [17, 19, 47]. However, we find that pelagic algae thrive in shallower lakes (*z*_m_ *≤* 10 m) regardless of the mixing depth, because these systems are too shallow for significant aphotic areas to form, thereby limiting the negative impact on downward mixing of pelagic algae in shallower lakes.

In addition, our simulations reveal that for deep Pattern 4 lakes (Fig. 2, *z*_m_ = 30 m), pelagic algae do not go extinct in large lakes under the same deep-mixing conditions that cause extinction in smaller lakes, all else being equal. This is because in large lakes, the relatively slow horizontal transport allows for the formation of nearshore refuges. These shallow, well-lit areas can sustain pelagic algal populations, sheltered from aphotic conditions at greater depths due to the limited horizontal transport, preventing the complete extinction seen in smaller deep lakes with a deep mixed layer. Moreover, we also find that a high nutrient content can counteract the extinction of pelagic algae when 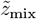 is large, due to increased growth of nutrient-limited pelagic algae in shallow areas (Fig. 2, *z*_m_ = 30, *N*_area_ *≥* 1000). Therefore, the detrimental effect of a deep mixing depth on pelagic algae in deep lakes is contingent on both lake size and nutrient content.

This result is particularly relevant as the critical depth principle was developed for deep oceanic systems [17, 19, 20], and our work demonstrates how it can be adapted to apply to freshwater lakes, where a combination of horizontal and vertical processes shapes algal distribution [21, 48–50].

Our model also confirms the unimodal relationship between pelagic biomass and 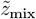, a pattern well-documented in both theoretical and empirical studies [15, 18, 23, 47]. This relationship arises from a trade-off: in very shallow mixed layers, nutrient supply is limited, while in very deep mixed layers, overall light conditions are too poor to support algal growth. The peak in biomass occurs at an intermediate mixing depth, where the upward transport of nutrients and the counteraction of downward algal sinking are optimally balanced against the detrimental effect of pelagic algae being mixed down to aphotic depths where algal growth is limited. The highest biomass is thus found at an intermediate 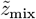, where a balance is achieved. Our study adds a new dimension to this concept by showing that this unimodal response is dependent on lake area. This suggests that the interplay between vertical mixing and horizontal mixing creates more complex dynamics than previously accounted for in one-dimensional models. We find that the fraction of total nutrients in pelagic algae changes in a complex, non-linear way with 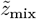 in deeper lakes (Fig. 7), and the specific shape of this response is different for lakes of different sizes.

### 5.2 New insights into benthic algal dynamics

Our study moves beyond the traditional focus on pelagic algae to provide novel insights into the role of 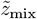 and lake size in benthic algal dynamics, a topic that to our knowledge has not been comprehensively addressed. We confirm established findings that greater lake depth and higher nutrient content negatively affect the contribution of benthic algae to total production [51–54]. Furthermore, we introduce a new finding: benthic algal biomass is also significantly influenced by 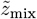 and lake size.

The relationship between benthic algal biomass and mixed layer depth is complex and dependent on lake depth. In shallow lakes (Fig. 7, *z*_m_ *≤* 3 m), benthic algal biomass shows either no response, or a negative response to initial deepening of the mixed layer. In contrast, in deep lakes (Fig. 7, *z*_m_ *≥*10 m), the pattern is the opposite: benthic algal biomass shows either no response, or increases at first as the mixed layer deepens, stabilizing for deeper mixing depths. This suggests that the effect of vertical mixing on benthic algae is mediated by a lake’s overall morphology. In shallow systems, a deeper mixed layer promotes the growth of pelagic algae by counteracting sinking and transporting nutrients upwards. Since light conditions at greater depths remain favorable, this results in enhanced pelagic algal growth which in turn increases competition at the detriment of benthic algae.

In deep lakes, the increased vertical mixing from a deep mixed layer efficiently transports nutrients from aphotic depths to well-lit nearshore areas, benefiting benthic algal growth. Crucially, this effect is not counteracted by an increase in pelagic algae as it is in shallow lakes, due to low pelagic algal growth at aphotic depths which allows the increased nutrient transport to result in a net benefit for the benthic algal community. Furthermore, our analysis reveals a new finding regarding the effect of lake size on benthic biomass. In shallow lakes, biomass either remains unchanged or increases with lake area. In contrast, in deep lakes, we observe the opposite pattern: benthic biomass decreases as lake area increases. An additional key finding from our simulations is the formation of a benthic deep chlorophyll maximum (DCM). Our model shows that this phenomenon, where benthic algal biomass peaks at an intermediate depth, can only arise if the mixed layer is shallower than the depth at which light limitation becomes dominant. This also requires that benthic algal growth is nutrient-limited in shallow areas and light-limited in deeper areas, which, along with a nutrient gradient that increases with depth, is necessary for the formation of a benthic deep chlorophyll maximum. This is clearly seen in our Pattern 2 simulations (Fig. 3, 4, panels on blue background), where a shallow thermocline allows for opposing vertical gradients of light and dissolved nutrients to form. This new phenomenon highlights how vertical stratification can be an important driver of the spatial heterogeneity of benthic algae.

### 5.3 Study limitations

Our modeling approach, while robust, has several limitations. First, we did not extensively analyze the dynamics of sedimented nutrients. This was a deliberate choice, as this is a quantity that is difficult to measure empirically, making it less suitable for model-data comparison. Second, we acknowledge that the background turbidity (*k*_bg_) significantly influences algal dynamics, as demonstrated in a related manuscript on unstratified lakes [38]. However, as our previous work found that varying *k*_bg_ did not affect the qualitative patterns and had a similar effect to varying the mean depth [38], we chose to focus on a single value for this parameter to simplify the analysis of the current study.

Finally, our fully factorial parameter sweep, while providing new insights into fundamental processes, includes some parameter combinations that are unlikely to occur in nature. For instance, mixing depth is often positively correlated with lake size [4, 14], while mean depth is typically negatively correlated with nutrient content [55, 56] and positively correlated with lake area [4, 57]. These correlations suggest that certain scenarios, such as deep mixed layers in small, shallow lakes or deep lakes with high nutrient concentrations, are less probable. However, we contend that our exhaustive exploration of parameter space is beneficial, as it more clearly reveals the complex dynamics and interactions between environmental drivers. Restricting our analysis to only the most plausible parameter combinations could obscure these underlying patterns, making it more difficult to fully understand the system’s behavior.

### 5.4 Outlook

Our results have implications for future research and regional upscaling. The strong similarity between the biomass patterns observed in this study and the qualitative phenomena found in an analogous model of unstratified lakes [38] suggests that at the whole-lake scale, both thermocline depth and lake size primarily affect algal dynamics by controlling the exchange rates of algae and nutrients between shallow and deep waters. This finding aligns with results from our previous work [37], which showed large differences in algal dynamics between lake2D and an analagous 1D model, further highlighting the critical importance of these exchange rates.

This highlights the potential of Lake2D as a tool for the regional upscaling of dynamics across ensembles of lakes, as demonstrated in a similar framework [58], where the use of 1D models that do not account for horizontal heterogeneity such as Global Lake Model (GLM), Arctic Lake Biogeochemistry Model (ALBM), PCLake+, and MyLake is currently the norm [59–62]. This approach could be further enhanced by incorporating the natural cross-correlations that exist between environmental drivers. By integrating these co-dependencies with our Lake2D model, future research can provide more realistic predictions of how climate change, which directly influences seasonal mixing regimes and the depth of the mixed layer in stratified lakes [63, 64], will affect algal dynamics in different types of lakes. Understanding these dynamics is more imperative now than ever.

## Appendix

### A Model description

In this appendix, we give a mathematical description of Lake2D, the model used in this work. We start in Section A.1 from a three-dimensional mathematical model of the dynamics of pelagic and benthic algae in a lake with arbitrary morphometry, and in Section A.2 we detail the mechanisms involved. In Section A.3, we derive Lake2D as a reduction of the three-dimensional mathematical model by assuming rotational symmetry of the problem.

#### A.1 Mathematical model

##### Lake domain and notation

Let the lake geometry be described by a three-dimensional domain Ω with piecewise smooth boundary *∂*Ω equipped with outward-pointing unit normal 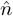. The domain of the lake bottom is denoted Γ and the domain of the lake surface is denoted *∂*Ω\ Γ. The lake surface *∂*Ω \Γ is horizontally flat at *z* = 0, where *z* denotes the depth below the surface. The boundary of the lake bottom, i.e., the lake shore, is denoted *∂*Γ and is equipped with outward pointing unit normal 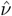(co-normal to Γ).

Below, spatial derivatives are written using coordinate-free vector calculus, where *∇* is the usual gradient operator in ℝ3 and 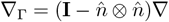 denotes the tangential gradient along Γ.

##### Governing equations

The model consists of the following five state variables: The concentrations of pelagic algal biomass *A* and a potentially growth-limiting inorganic nutrient *N*_d_ in the lake water, the areal densities of sedimented particular nutrient *N*_s_ at the lake bottom and of benthic algal biomass *B* at the sediment-water interface, and the intensity of the potentially growth-limiting light *I* in the water column. Benthic and pelagic algal biomass is expressed in units of carbon, and the dissolved and sedimented nutrient is assumed to be phosphorus. The rates of change of *{A, N*_d_, *N*_s_, *B}* are governed by the following system of differential equations

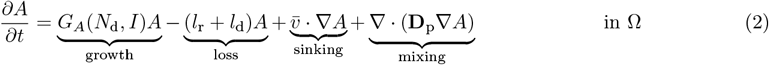

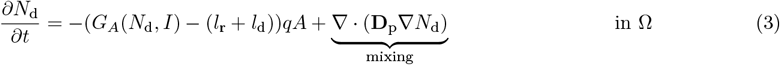

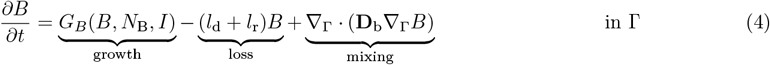

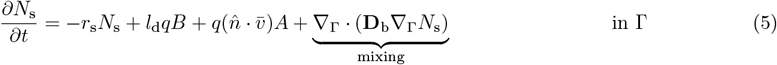

with boundary conditions for the bulk fields *{A, N*_d_*}*

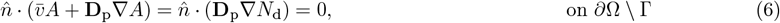

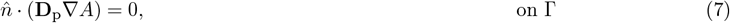

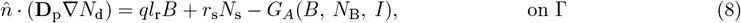

and boundary conditions for the bottom fields *{B, N*_s_*}*

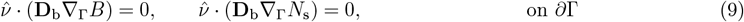

The last state variable, the local light intensity *I* at a depth *z* below the lake surface, is given by

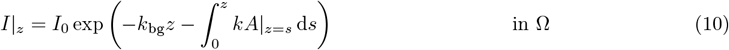

The growth rates of pelagic and benthic algae, *G*_*A*_, and *G*_*B*_, are given by

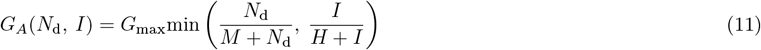

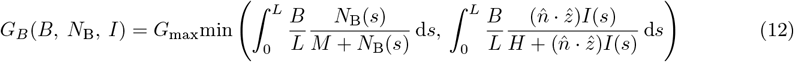

where 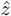 is a unit vector pointing downwards.

All variables and parameters are listed with their units in Table 1, using values based on previous work [13, 18, 52]. These governing equations are explained in detail in the next section.

#### A.2 Description of governing equations

##### Transport mechanisms

The model considers two spatial transport processes; turbulent mixing and sinking. Turbulent mixing of pelagic algae *A* and dissolved nutrients *N*_d_ is described by vertical and horizontal diffusion. In addition, benthic algae and sedimented nutrients experience diffusion along the bottom (typically very slow). The amount of diffusion in different spatial directions, in the water body Ω respectively along the the bottom Γ, is described through the matrices **D**_p_ respectively **D**_b_, which contain the diffusion coefficients. These coefficients can depend on the spatial position (specifically on the depth *z* in this work). Sinking of pelagic algae is described as a transport term 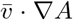 in (2), where pelagic algae is transported at a constant velocity 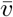 (a vector pointing vertically downwards). Pelagic algae that reach the bottom die and their nutrients turn into sedimented particular nutrients. Since we assume that the system is closed for nutrients, the zero-flux boundary conditions in (6), (7) and (9) follow.

##### Pelagic algal dynamics

The growth of pelagic algae at a given position in the lake is either limited by the local light intensity, or the local dissolved nutrient concentration, as described by the minimum function of two Monod terms in (11) where *G*_max_ is the maximum specific growth rate and *H* and *M* are the half-saturation constants of light and nutrient-limited growth, respectively. Light intensity *I* attenuates vertically in the water column according to (10), where pelagic algae contribute with specific attenuation coefficient *k* and background attenuation with coefficient *k*_bg_. The latter represents light attenuation by non-algal materials in the water and by water itself. In addition to sedimentation losses, pelagic biomass is lost through background respiration and mortality with fixed rates *l*_r_ and *l*_d_, respectively.

##### Pelagic nutrient dynamics

Uptake and recycling of dissolved nutrients at a given depth are driven by local algal growth and losses. We assume that the stoichiometric phosphorus-to-carbon (P:C) ratio *q* of algal biomass is fixed. Consequently, the local uptake of dissolved phosphorus equals a fraction *q* of algal carbon production. Similarly, algal (carbon) biomass losses through mortality and respiration are accompanied by proportional phosphorus losses, which we assume to be instantly recycled in dissolved form.

##### Sediment nutrient dynamics

Particular nutrients enter the sediment through two processes; sinking of pelagic algae and background mortality of benthic algae at the rate *l*_d_. We assume that remineralization of sedimented nutrients is a first order process at the rate *r*_s_ and that remineralized nutrients are released into the water directly above the bottom, see (8).

##### Benthic algal dynamics

To focus our analyses on the impact of physical factors, we assume that benthic and pelagic algae have identical traits for growth and metabolism. Thus, benthic and pelagic algae have identical maximum specific growth rates and half saturation constants for light- and nutrient-dependent growth, identical light attenuation coefficients and elemental P:C ratios, as well as identical respiration and mortality rates.

To calculate their growth rate, we assume that benthic algae form a homogeneous layer of thickness *L*, which depends on benthic algal biomass as

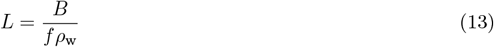

where *f* is the ratio of benthic algal carbon to wet mass and *ρ*_w_ is the specific density of algal wet mass assumed to be identical to the density of water. Light in the benthic algal layer is attenuated according to Lambert-Beer’s law. Assuming that non-algal light attenuation is negligible, light-limited benthic algal growth can then be integrated over the benthic algal layer as [65]

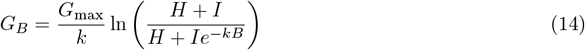

where *I* is the local light intensity at the bottom of the water column.

Nutrient-limited benthic algal growth is modeled analogously by integrating over the vertical profile of dissolved nutrients in the benthic algal layer. We assume that dissolved nutrients immediately above the lake bottom diffuse into the benthic algal layer while simultaneously being consumed by benthic algae. These two processes are modeled with the differential equation

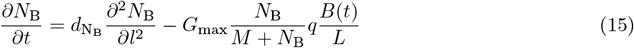

where *N*_B_ is the concentration of dissolved nutrient in the benthic algal layer, 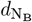 is the diffusion coefficient in the benthic algal layer, *l ∈* [0, *L*] denotes the orthogonal position inside the benthic algal layer, and *q* is the P:C ratio of benthic biomass. We assume that these processes are fast compared to other processes, and therefore treat (15) on a separate timescale. For each time *t*, we approximate the steady state solution of (15) by linearizing the benthic algal consumption term and solving the resulting equation analytically. We further assume that the nutrient concentration at the surface of the benthic algal layer equals the concentration at the bottom of the water column and that nutrients do not diffuse out of the bottom of the benthic algal layer, yielding the boundary conditions

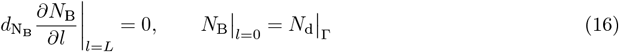

where Γ is the lake bottom and 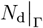 is the dissolved nutrient concentration along the bottom of the lake. We then integrate the nutrient-limited growth over the benthic algal layer (first integral in (12)), using the approximated solution of (15) for each time step. In a final step, benthic algal growth per unit area is calculated as the minimum of nutrient and light-dependent growth (12). This approach can sometimes overestimate benthic algal growth, since it is possible that growth switches from nutrient to light limitation somewhere inside the benthic algal layer. Note that *N*_B_ is a dummy variable that is only used to calculate nutrient-limited benthic algal growth and therefore does not contribute to the nutrient mass balance of the system.

##### Nutrient fluxes at the sediment-water interface

We assume that nutrients lost through benthic algal respiration are released in dissolved form into the water directly above, which is the same water from which the nutrients are drawn that sustain benthic algal growth. Together with the release of remineralized particular nutrients from the sediment, this yields the boundary condition for dissolved nutrients at the bottom of the water column (8).

#### A.2.1 Nutrient conservation

The total nutrient content of a modeled lake is defined as total phosphorus per unit surface area

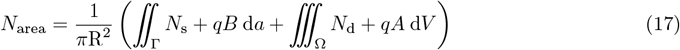

where Γ is the bottom surface, and Ω is the lake volume.The absence of internal nutrient sources and the closed boundary conditions imply that *N*_area_ is a measure of nutrient enrichment, which is determined by the initial conditions and remains constant over time.

#### A.3 Reduced model - Lake2D

Lake2D is derived from the three-dimensional mathematical model described in Section A.1 by assuming radial symmetry. This means both a radially symmetric lake geometry as well as radial symmetry of the state variable fields *{A, N*_d_, *N*_s_, *B, I}*. In cylindrical coordinates *{r, φ, z}*, where *r* is the horizontal distance to the lake center axis, *φ* is the azimuthal direction, and *z*, as before, is the depth below the surface, the assumption of radial symmetry means all spatially dependent terms are constant with respect to *φ*. Since all derivatives with respect to *φ* will be zero, this will simplify the governing equations, and any integrals over Ω or Γ can be reduced to two-dimensional integrals over a radial cross section of Ω or Γ by integrating away the azimuthal direction. Hence, the model is effectively reduced to a problem of two spatial dimensions *{r, z}*, where it is sufficient to track variables on a radial cross-section of the lake geometry. This approach implies a reflective boundary condition for pelagic algae and dissolved nutrients along the left boundary of the cross-section, corresponding to the center of the lake

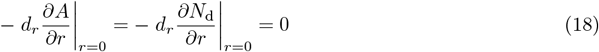

and correspondingly for the fields along the bottom we have

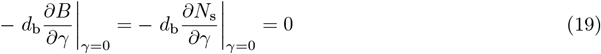

where *γ* is the arc length along the radial cross-section of Γ with *γ* = 0 at the center.

##### Implementation aspects

To facilitate execution and analysis, we non-dimensionalized the model, see [38, Appendix A], and implemented it on a radial cross-section of the lake. In this and previous works [37, 38], we have chosen to use a cone as our radially symmetric lake geometry, since its simple topography facilitates visualization and interpretation of results, and also because its bottom profile is similar to the hypsography of many real lakes [39–41].

## References

[1] S. R. Carpenter, “Lake geometry: implications for production and sediment accretion rates,” Journal of Theoretical Biology, vol. 105, no. 2, pp. 273–286, 1983.

[2] M.-E. Ferland, Y. T. Prairie, C. Teodoru, and P. A. del Giorgio, “Linking organic carbon sedimentation, burial efficiency, and long-term accumulation in boreal lakes,” Journal of Geophysical Research: Biogeosciences, vol. 119, no. 5, pp. 836–847, 2014.

[3] Y. Vadeboncoeur, G. Peterson, M. J. Vander Zanden, and J. Kalff, “Benthic algal production across lake size gradients: interactions among morphometry, nutrients, and light,” Ecology, vol. 89, no. 9, pp. 2542–2552, 2008.

[4] E. Fee, R. Hecky, S. Kasian, and D. Cruikshank, “Effects of lake size, water clarity, and climatic variability on mixing depths in Canadian Shield lakes,” Limnology and oceanography, vol. 41, no. 5, pp. 912–920, 1996.

[5] D. M. Imboden and A. Wüest, “Mixing mechanisms in lakes,” in Physics and chemistry of lakes, pp. 83–138, Springer, 1995.

[6] L. Håkanson, “The importance of lake morphometry for the structureand function of lakes,” International Review of Hydrobiology: A Journal Covering all Aspects of Limnology and Marine Biology, vol. 90, no. 4, pp. 433–461, 2005.

[7] P. Chow-Fraser, “Use of the morphoedaphic index to predict nutrient status and algal biomass in some Canadian lakes,” Canadian Journal of Fisheries and Aquatic Sciences, vol. 48, p. 1909–1918, Oct. 1991.

[8] L. Håkanson, Lakes: form and function. Blackburn Press Caldwell, New Jersey, 2004.

[9] B. Nixdorf and R. Deneke, “Why ‘very shallow’ lakes are more successful opposing reduced nutrient loads,” Hydrobiologia, vol. 342–343, p. 269–284, Jan. 1997.

[10] B. Boehrer and M. Schultze, “Stratification of lakes,” Reviews of Geophysics, vol. 46, no. 2, 2008.

[11] W. M. Lewis Jr, “A revised classification of lakes based on mixing,” Canadian Journal of Fisheries and Aquatic Sciences, vol. 40, no. 10, pp. 1779–1787, 1983.

[12] R. I. Woolway and C. J. Merchant, “Worldwide alteration of lake mixing regimes in response to climate change,” Nature Geoscience, vol. 12, no. 4, pp. 271–276, 2019.

[13] P. T. Kelly, C. T. Solomon, J. A. Zwart, and S. E. Jones, “A framework for understanding variation in pelagic gross primary production of lake ecosystems,” Ecosystems, vol. 21, pp. 1364–1376, 2018.

[14] P. D. Isles, I. F. Creed, A. Jonsson, and A.-K. Bergström, “Trade-offs between light and nutrient availability across gradients of dissolved organic carbon lead to spatially and temporally variable responses of lake phytoplankton biomass to browning,” Ecosystems, vol. 24, no. 8, pp. 1837–1852, 2021.

[15] J. Huisman and F. J. Weissing, “Competition for nutrients and light in a mixed water column: a theoretical analysis,” The American Naturalist, vol. 146, no. 4, pp. 536–564, 1995.

[16] R. Ptacnik, A. G. Solimini, T. Andersen, T. Tamminen, P. Brettum, L. Lepistö, E. Willén, and S. Rekolainen, “Diversity predicts stability and resource use efficiency in natural phytoplankton communities,” Proceedings of the National Academy of Sciences, vol. 105, no. 13, pp. 5134–5138, 2008.

[17] J. Huisman, P. van Oostveen, and F. J. Weissing, “Critical depth and critical turbulence: two different mechanisms for the development of phytoplankton blooms,” Limnology and oceanography, vol. 44, no. 7, pp. 1781–1787, 1999.

[18] C. G. Jäger, S. Diehl, and M. Emans, “Physical determinants of phytoplankton production, algal stoichiometry, and vertical nutrient fluxes,” The American Naturalist, vol. 175, no. 4, pp. E91–E104, 2010.

[19] H. U. Sverdrup, “On conditions for the vernal blooming of phytoplankton,” J. Cons. Int. Explor. Mer, vol. 18, no. 3, pp. 287–295, 1953.

[20] D. Siegel, S. Doney, and J. Yoder, “The North Atlantic spring phytoplankton bloom and Sverdrup’s critical depth hypothesis,” science, vol. 296, no. 5568, pp. 730–733, 2002.

[21] S. Diehl, S. A. Berger, Q. Soissons, D. P. Giling, and H. Stibor, “An experimental demonstration of the critical depth principle,” ICES Journal of Marine Science, vol. 72, no. 6, pp. 2051–2060, 2015.

[22] R. Ptacnik, S. Diehl, and S. Berger, “Performance of sinking and nonsinking phytoplankton taxa in a gradient of mixing depths,” Limnology and Oceanography, vol. 48, no. 5, pp. 1903–1912, 2003.

[23] S. A. Berger, S. Diehl, T. J. Kunz, D. Albrecht, A. M. Oucible, and S. Ritzer, “Light supply, plankton biomass, and seston stoichiometry in a gradient of lake mixing depths,” Limnology and oceanography, vol. 51, no. 4, pp. 1898–1905, 2006.

[24] S. Diehl, S. Berger, R. Ptacnik, and A. Wild, “Phytoplankton, light, and nutrients in a gradient of mixing depths: field experiments,” Ecology, vol. 83, no. 2, pp. 399–411, 2002.

[25] J. B. Butcher, D. Nover, T. E. Johnson, and C. M. Clark, “Sensitivity of lake thermal and mixing dynamics to climate change,” Climatic Change, vol. 129, no. 1, pp. 295–305, 2015.

[26] R. P. North, R. L. North, D. M. Livingstone, O. Köster, and R. Kipfer, “Long-term changes in hypoxia and soluble reactive phosphorus in the hypolimnion of a large temperate lake: consequences of a climate regime shift,” Global change biology, vol. 20, no. 3, pp. 811–823, 2014.

[27] Y. Yankova, S. Neuenschwander, O. Köster, and T. Posch, “Abrupt stop of deep water turnover with lake warming: Drastic consequences for algal primary producers,” Scientific Reports, vol. 7, no. 1, p. 13770, 2017.

[28] M. Cantonati and R. L. Lowe, “Lake benthic algae: toward an understanding of their ecology,” Freshwater Science, vol. 33, no. 2, pp. 475–486, 2014.

[29] I. C. Puts, J. Ask, M. B. Siewert, R. A. Sponseller, D. O. Hessen, and A.-K. Bergström, “Landscape determinants of pelagic and benthic primary production in northern lakes,” Global Change Biology, vol. 28, no. 23, pp. 7063–7077, 2022.

[30] Y. Vadeboncoeur and A. D. Steinman, “Periphyton function in lake ecosystems,” The Scientific World Journal, vol. 2, no. 1, pp. 1449–1468, 2002.

[31] D. M. DeNicola and M. Kelly, “Role of periphyton in ecological assessment of lakes,” Freshwater Science, vol. 33, no. 2, pp. 619–638, 2014.

[32] C. S. Rousseaux and W. W. Gregg, “Interannual variation in phytoplankton primary production at a global scale,” Remote sensing, vol. 6, no. 1, pp. 1–19, 2013.

[33] J. Ask, J. Karlsson, L. Persson, P. Ask, P. Byström, and M. Jansson, “Whole-lake estimates of carbon flux through algae and bacteria in benthic and pelagic habitats of clear-water lakes,” Ecology, vol. 90, no. 7, pp. 1923–1932, 2009.

[34] J. Karlsson, P. Byström, J. Ask, P. Ask, L. Persson, and M. Jansson, “Light limitation of nutrient-poor lake ecosystems,” Nature, vol. 460, no. 7254, pp. 506–509, 2009.

[35] I. Puts, A.-K. Bergström, H. Verheijen, S. Norman, and J. Ask, “An ecological and methodological assessment of benthic gross primary production in northern lakes,” Ecosphere, vol. 13, no. 3, p. e3973, 2022.

[36] Y. Vadeboncoeur, D. M. Lodge, and S. R. Carpenter, “Whole-lake fertilization effects on distribution of primary production between benthic and pelagic habitats,” Ecology, vol. 82, no. 4, pp. 1065–1077, 2001.

[37] H. Harlin, K. Larsson, and S. Diehl, “Can whole-lake algal biomass be captured by one-dimensional modeling approaches? An exploration using ‘Lake2D’,” BioRxiv, 2025. [Preprint].

[38] H. Harlin, K. Larsson, Å. Brännström, and S. Diehl, “Physical drivers of benthic and pelagic algal biomass dynamics in lakes: a conceptual exploration with ‘Lake2D’,” BioRxiv, 2025. [Preprint].

[39] L. Håkanson, “On lake form, lake volume and lake hypsographic survey,” Geografiska Annaler: Series A, Physical Geography, vol. 59, no. 1-2, pp. 1–29, 1977.

[40] J. Stachelek, P. J. Hanly, and P. A. Soranno, “Imperfect slope measurements drive overestimation in a geometric cone model of lake and reservoir depth,” Inland Waters, vol. 12, no. 2, pp. 283–293, 2022.

[41] M. Klaus, H. A. Verheijen, J. Karlsson, and D. A. Seekell, “Depth and basin shape constrain ecosystem metabolism in lakes dominated by benthic primary producers,” Limnology and Oceanography, vol. 67, no. 12, pp. 2763–2778, 2022.

[42] D. Seekell and B. Cael, “Why does the relationship between benthic primary production and lake morphometry vary regionally?,” Aquatic Sciences, vol. 85, no. 3, p. 87, 2023.

[43] B. Cael, A. Heathcote, and D. Seekell, “The volume and mean depth of Earth’s lakes,” Geophysical Research Letters, vol. 44, no. 1, pp. 209–218, 2017.

[44] M. Chen, G. Zeng, J. Zhang, P. Xu, A. Chen, and L. Lu, “Global landscape of total organic carbon, nitrogen and phosphorus in lake water,” Scientific reports, vol. 5, no. 1, p. 15043, 2015.

[45] S. Sobek, L. J. Tranvik, Y. T. Prairie, P. Kortelainen, and J. J. Cole, “Patterns and regulation of dissolved organic carbon: An analysis of 7,500 widely distributed lakes,” Limnology and oceanography, vol. 52, no. 3, pp. 1208–1219, 2007.

[46] C. Verpoorter, T. Kutser, D. A. Seekell, and L. J. Tranvik, “A global inventory of lakes based on high-resolution satellite imagery,” Geophysical Research Letters, vol. 41, no. 18, pp. 6396–6402, 2014.

[47] S. Diehl, “Phytoplankton, light, and nutrients in a gradient of mixing depths: theory,” Ecology, vol. 83, no. 2, pp. 386–398, 2002.

[48] F. Peeters, D. Straile, A. Lorke, and D. Ollinger, “Turbulent mixing and phytoplankton spring bloom development in a deep lake,” Limnology and oceanography, vol. 52, no. 1, pp. 286–298, 2007.

[49] E. Gronchi, K. D. Jöhnk, D. Straile, S. Diehl, and F. Peeters, “Local and continental-scale controls of the onset of spring phytoplankton blooms: Conclusions from a proxy-based model,” Global Change Biology, vol. 27, no. 9, pp. 1976–1990, 2021.

[50] S. MacIntyre and J. M. Melack, “Vertical and horizontal transport in lakes: Linking littoral, benthic, and pelagic habitats,” Journal of the North American Benthological Society, vol. 14, p. 599–615, Dec. 1995.

[51] C. G. Jäger and S. Diehl, “Resource competition across habitat boundaries: asymmetric interactions between benthic and pelagic producers,” Ecological Monographs, vol. 84, no. 2, pp. 287–302, 2014.

[52] F. Rivera Vasconcelos, S. Diehl, P. Rodríguez, J. Karlsson, and P. Byström, “Effects of terrestrial organic matter on aquatic primary production as mediated by pelagic–benthic resource fluxes,” Ecosystems, vol. 21, pp. 1255–1268, 2018.

[53] L.-A. Hansson, “Factors regulating periphytic algal biomass,” Limnology and Oceanography, vol. 37, no. 2, pp. 322–328, 1992.

[54] Y. Vadeboncoeur, E. Jeppesen, M. J. V. Zanden, H.-H. Schierup, K. Christoffersen, and D. M. Lodge, “From Greenland to green lakes: cultural eutrophication and the loss of benthic pathways in lakes,” Limnology and oceanography, vol. 48, no. 4, pp. 1408–1418, 2003.

[55] P. A. Staehr, L. Baastrup-Spohr, K. Sand-Jensen, and C. Stedmon, “Lake metabolism scales with lake morphometry and catchment conditions,” Aquatic Sciences, vol. 74, pp. 155–169, 2012.

[56] T. Noges, “Relationships between morphometry, geographic location and water quality parameters of European lakes,” Hydrobiologia, vol. 633, no. 1, pp. 33–43, 2009.

[57] C. K. Minns, J. E. Moore, B. J. Shuter, and N. E. Mandrak, “A preliminary national analysis of some key characteristics of Canadian lakes,” Canadian Journal of Fisheries and Aquatic Sciences, vol. 65, no. 8, pp. 1763–1778, 2008.

[58] C. Gudasz, D. Vachon, and Y. T. Prairie, “A comprehensive framework for integrating lake hypsography and function on a global scale,” Nature Water, vol. 3, no. 7, pp. 818–830, 2025.

[59] M. Golub, W. Thiery, R. Marcé, D. Pierson, I. Vanderkelen, D. Mercado, R. I. Woolway, L. Grant, E. Jennings, J. Schewe, et al., “A framework for ensemble modelling of climate change impacts on lakes worldwide: the ISIMIP Lake Sector,” Geoscientific Model Development Discussions, vol. 2022, pp. 1–57, 2022.

[60] L. Råman Vinnå, I. Medhaug, M. Schmid, and D. Bouffard, “The vulnerability of lakes to climate change along an altitudinal gradient,” Communications Earth & Environment, vol. 2, no. 1, p. 35, 2021.

[61] M. R. Hipsey, L. C. Bruce, C. Boon, B. Busch, C. C. Carey, D. P. Hamilton, P. C. Hanson, J. S. Read, E. De Sousa, M. Weber, et al., “A General Lake Model (GLM 3.0) for linking with high-frequency sensor data from the Global Lake Ecological Observatory Network (GLEON),” Geoscientific Model Development, vol. 12, no. 1, pp. 473–523, 2019.

[62] L. Kramer, T. A. Troost, A. B. Janssen, R. J. Brederveld, L. P. van Gerven, D. van Wijk, W. M. Mooij, and S. Teurlincx, “Connecting lakes: Modeling flows and interactions of organisms and matter throughout the waterscape,” Environmental Modelling & Software, vol. 167, p. 105765, 2023.

[63] B. M. Kraemer, O. Anneville, S. Chandra, M. Dix, E. Kuusisto, D. M. Livingstone, A. Rimmer, S. G. Schladow, E. Silow, L. M. Sitoki, et al., “Morphometry and average temperature affect lake stratification responses to climate change,” Geophysical Research Letters, vol. 42, no. 12, pp. 4981–4988, 2015.

[64] T. Shatwell, W. Thiery, and G. Kirillin, “Future projections of temperature and mixing regime of European temperate lakes,” Hydrology and Earth System Sciences, vol. 23, no. 3, pp. 1533–1551, 2019.

[65] J. Huisman and F. J. Weissing, “Light-limited growth and competition for light in well-mixed aquatic environments: an elementary model,” Ecology, vol. 75, no. 2, pp. 507–520, 1994.

[66] F. Peeters and H. Hofmann, “Length-scale dependence of horizontal dispersion in the surface water of lakes,” Limnology and Oceanography, vol. 60, no. 6, pp. 1917–1934, 2015.

[67] C. Murthy, “Horizontal diffusion characteristics in Lake Ontario,” Journal of physical oceanography, vol. 6, no. 1, pp. 76–84, 1976.

[68] J. Huisman, J. Sharples, J. M. Stroom, P. M. Visser, W. E. A. Kardinaal, J. M. Verspagen, and B. Sommeijer, “Changes in turbulent mixing shift competition for light between phytoplankton species,” Ecology, vol. 85, no. 11, pp. 2960–2970, 2004.

[69] B. Boehrer, J. Ilmberger, and K. O. Münnich, “Vertical structure of currents in western Lake Constance,” Journal of Geophysical Research: Oceans, vol. 105, no. C12, pp. 28823–28835, 2000.

[70] F. Peeters, A. Wüest, G. Piepke, and D. M. Imboden, “Horizontal mixing in lakes,” Journal of Geophysical Research: Oceans, vol. 101, no. C8, pp. 18361–18375, 1996.

[71] D. Imboden and S. Emerson, “Natural radon and phosphorus as limnologic tracers: Horizontal and vertical eddy diffusion in Greifensee,” Limnology and Oceanography, vol. 23, no. 1, pp. 77–90, 1978.

[72] J. Huisman, N. N. Pham Thi, D. M. Karl, and B. Sommeijer, “Reduced mixing generates oscillations and chaos in the oceanic deep chlorophyll maximum,” Nature, vol. 439, no. 7074, pp. 322–325, 2006.

